# A decade after being listed as Endangered: Japanese eel stock inferred from fishery-dependent and independent monitoring records (Preprint)

**DOI:** 10.1101/2025.08.05.668826

**Authors:** Kenzo Kaifu, Hikaru Itakura, Tomonari Kotani, Akira Shinoda, Yu-San Han, Tatsuki Yoshinaga, Ryoshiro Wakiya

## Abstract

This study assesses recent trends in the abundance of Japanese eels (*Anguilla japonica*), which have been listed as Endangered on the IUCN Red List of Threatened Species since 2014, by updating previously reported coastal fisheries datasets and incorporating new freshwater data and scientific monitoring records for glass eels in Japan and Taiwan. Catch-per-unit-effort (CPUE) data for yellow and silver eels revealed statistically significant declines in seven of eight datasets, with projected reductions over three generations (24 years) ranging from 79.2% to 99.9%. In contrast, no significant temporal trends were detected in the CPUE of glass eels, likely due to high interannual variability driven by oceanic conditions. The integration of freshwater and coastal data, along with fishery-independent monitoring, enhances the reliability of these indicators. The results provided in this study represent the best currently available indicators of Japanese eel population dynamics.

## Introduction

The Japanese eel, *Anguilla japonica*, is a catadromous fish species that spawns in waters west of the Mariana Islands (Tsukamoto et al. 2011). Their leaf-like leptocephalus larvae drift with ocean currents before migrating to freshwater and estuarine habitats, where they metamorphose into ‘glass eels’. These eels spend most of their lives as ‘yellow eels’ in continental waters for varying durations before undergoing puberty. Once entering puberty, they metamorphose into ‘silver eels’ and return to the open ocean to spawn, after which they are thought to die, exhibiting a semelparous reproductive strategy (Aoyama and Miller 2003). In East Asia, this species is highly commercially important and recruiting glass eels are intensively captured for use in aquaculture (Shiraishi and Crook, 2015). However, despite its high commercial value and decades of research into aquaculture technologies, the artificial propagation of this species remains confined to laboratory settings and has not yet been achieved on a commercially viable scale, and commercial production still relies entirely on wild-caught glass eels (Righton et al. 2021). The Japanese eel population has decreased since the 1970s and they are now in a historically critical situation, with the species having been listed as Endangered (EN) on the IUCN Red List of Threatened Species since 2014 (Pike et al., 2020a).

Due to its catadromous life cycle, which involves long-distance migrations between freshwater and marine environments, studying the population dynamics of anguillid eels presents considerable challenges. A pioneering attempt to model the population dynamics of the Japanese eel was made by Tanaka (2014), who developed a stock assessment model despite the severe limitations in ecological and fishery data for this species. Although the Japanese eel is distributed throughout East Asia, Tanaka’s age- and sex-structured production model relied exclusively on Japanese fisheries data, supplementing ecological parameters such as mortality rates with information derived from other anguillid species, particularly the European eel (Tanaka 2014).

While subsequent research may enable more robust modelling as data availability improves, the uncertainty currently surrounding stock assessments for the Japanese eel remains substantial and should not be used to guide management policy (Kaifu 2019). In such cases, where detailed stock assessment is impractical, simple indicators such as catch-per-unit-effort (CPUE) can serve as useful proxies for monitoring population trends. Kaifu and Yokouchi (2019) analysed CPUE data from six coastal fisheries in Japan, collected through a nationwide survey conducted by the Fisheries Agency. That study revealed statistically significant declines in CPUE for four of the six fisheries, with no significant trends detected in the remaining two. On this basis, it was concluded that Japanese eel abundance was declining in Japan’s coastal waters (Kaifu and Yokouchi 2019). However, the study had several limitations. First, the period of the two datasets without significant trends were 8 years, which was very short and may have hindered the detection of any actual temporal trends. Second, the study was restricted to coastal areas and did not include freshwater fisheries intentionally due to the confounding effect of eel restocking, which is widespread in Japanese freshwater systems and complicates assessments of natural recruitment (Kaifu et al. 2018; Itakura et al. 2019; Kaifu 2019). Finally, although the study examined fisheries data for glass eels in Japan, these datasets likely included a substantial proportion of unreported catches, making them unsuitable for assessing population trends (Kaifu 2019).

The present study aims to update and expand upon the previous work of Kaifu and Yokouchi (2019) by addressing these three key limitations and providing a clearer understanding of population trends in the Japanese eel, one of the most commercially important species in East Asia. First, the six coastal datasets used in the 2019 study were updated and re-analysed. Second, new datasets were included from freshwater fisheries and scientific monitoring efforts, thereby incorporating data from inland as well as coastal areas. It should be noted that, in these freshwater sites, the proportion of stocked eels was confirmed to be low, reducing the likelihood of confounding effects. Lastly, for glass eels, we analysed scientific monitoring data collected in Japan and Taiwan, including data obtained through collaborations between researchers and fishers, thereby minimising the potential influence of unreported catch. This study contributes to a better understanding of the population dynamics of this economically valuable species by providing multiple abundance indices derived from diverse and more robust data sources.

## Materials and Methods

### Data Collection

Figure 1 indicates the areas from which data were obtained for this study, and details of all datasets analysed in this study are summarised in Table 1. For yellow and silver eels, updated fisheries data were obtained from the same fisheries cooperative associations or individual fishers surveyed by Kaifu and Yokouchi (2019), extending the dataset up to the year 2024. In some cases, where the original fisheries were no longer active, data were available only up to 2022. Detailed descriptions of the environments of these fishing grounds are provided in Kaifu and Yokouchi (2019), but most of them are located in shallow coastal areas, typically less than 10 m in depth. To ensure the confidentiality of all personal information, the names of the fisheries cooperative associations and individual fishers who provided fisheries data are not disclosed. Only CPUE values are presented to avoid revealing actual catch amounts, following the approach of Kaifu and Yokouchi (2019).

**Table 1.**
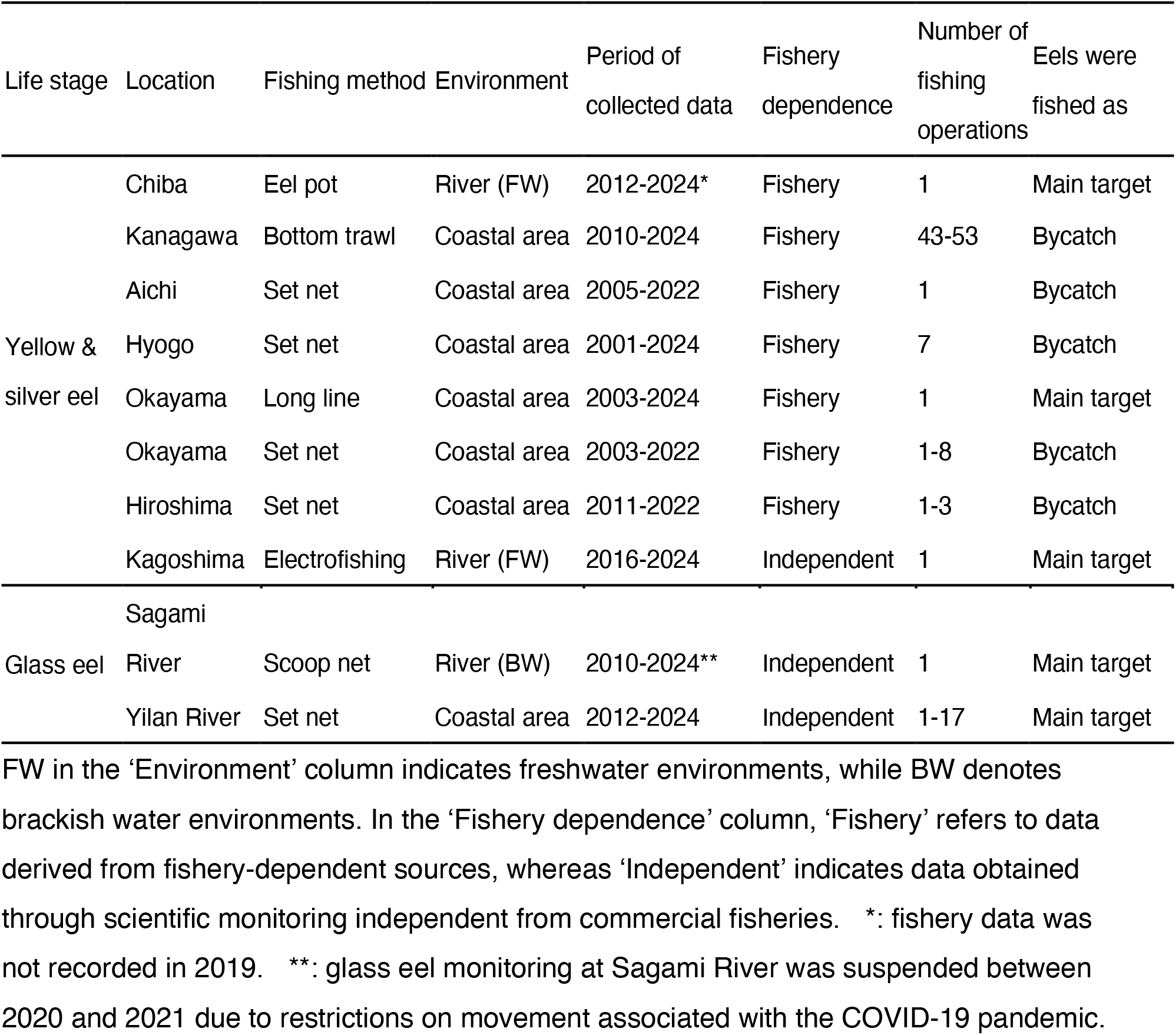
Data sets used in this study.

**Fig. 1.**
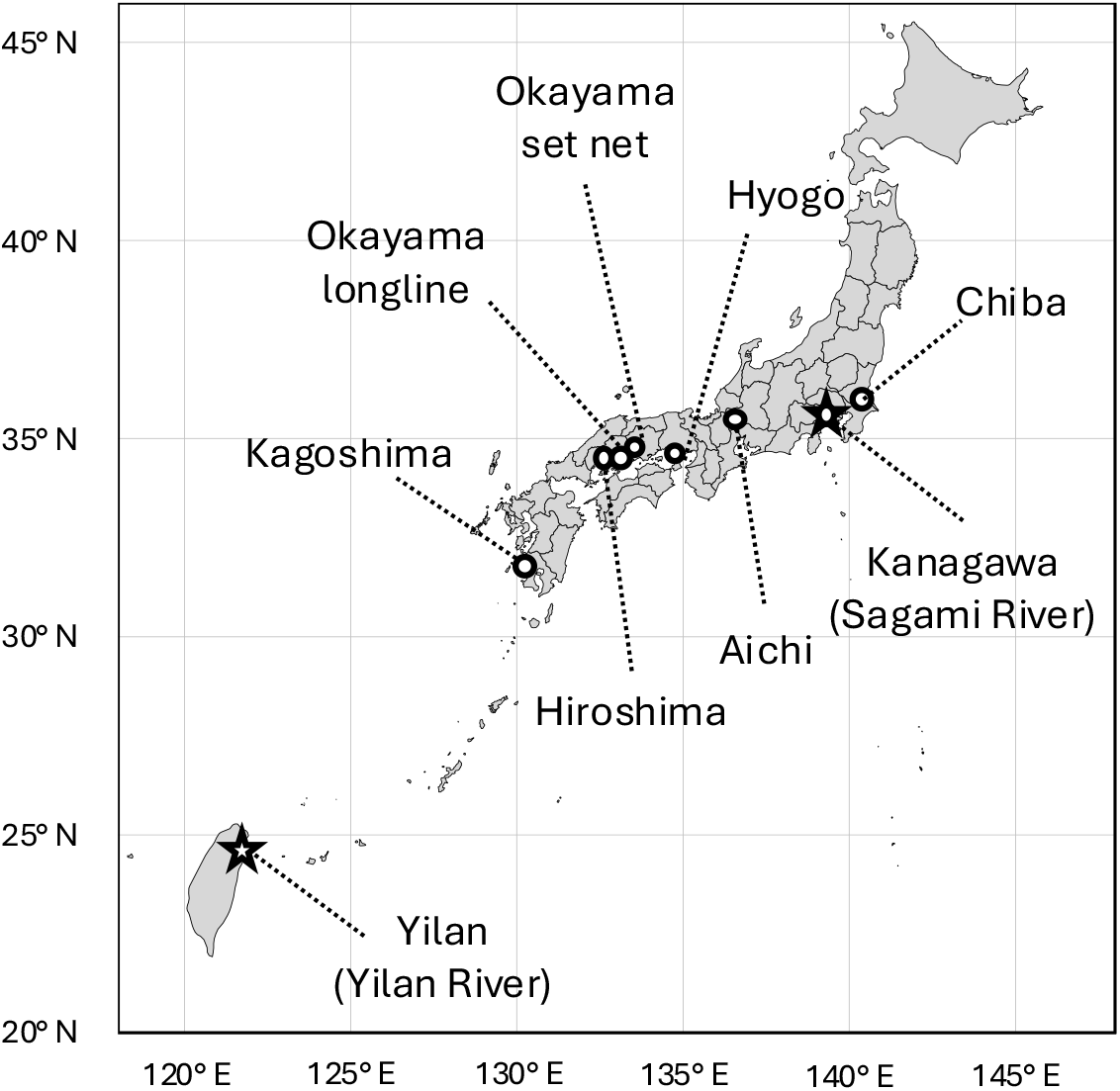
Locations where eel abundance data were obtained Circles indicate areas where data on yellow and silver eels were collected, while stars indicate areas where data on glass eels were obtained. Kanagawa is a region where fishery data for yellow and silver eels were collected, and it is also the location of the Sagami River, where glass eel monitoring was conducted. For details of the data collected, see Table 1.

In addition to these coastal datasets, freshwater data were also analysed, including records from an eel fishery in Chiba and a scientific monitoring programme in Kagoshima. For Chiba, fisheries data was obtained from a fisher who has fished eels using eel pots in freshwater area of the lower Tone River. In this area, no eel stocking has been conducted recently, and Itakura et al. (2019) estimated that 97% of eels inhabiting this area consists of naturally recruited wild eels, based on otolith stable isotopes. The Tone River has the second longest stream length (322 km) and the largest catchment area (16,840 km^2^) in Japan. For more detailed information, see Figure 2 in Itakura et al. (2019). Sampling site C on that map corresponds to the water body from which the fishery data used in this study were obtained. From Kagoshima, we have annually conducted scientific monitoring since 2016 in the Kaizoko River using electrofishing. Kaizoko River is a small river with only 3.3 km length. Because no commercial eel fishery has been conducted, no eel have been stocked into the river by fishers. Although eels have been released for research purposes a few times, all released individuals were marked with PIT tags, allowing clear distinction from naturally recruited individuals (Wakiya et al. 2022). This study analysed only naturally recruited individuals. Ten study sites of 40 m length were set up in the river, and subsequent quantitative electrofishing surveys were conducted annually at the same study sites until 2024. For more details such as biotic and abiotic environment information, please see Wakiya et al. (2022).

**Fig. 2.**
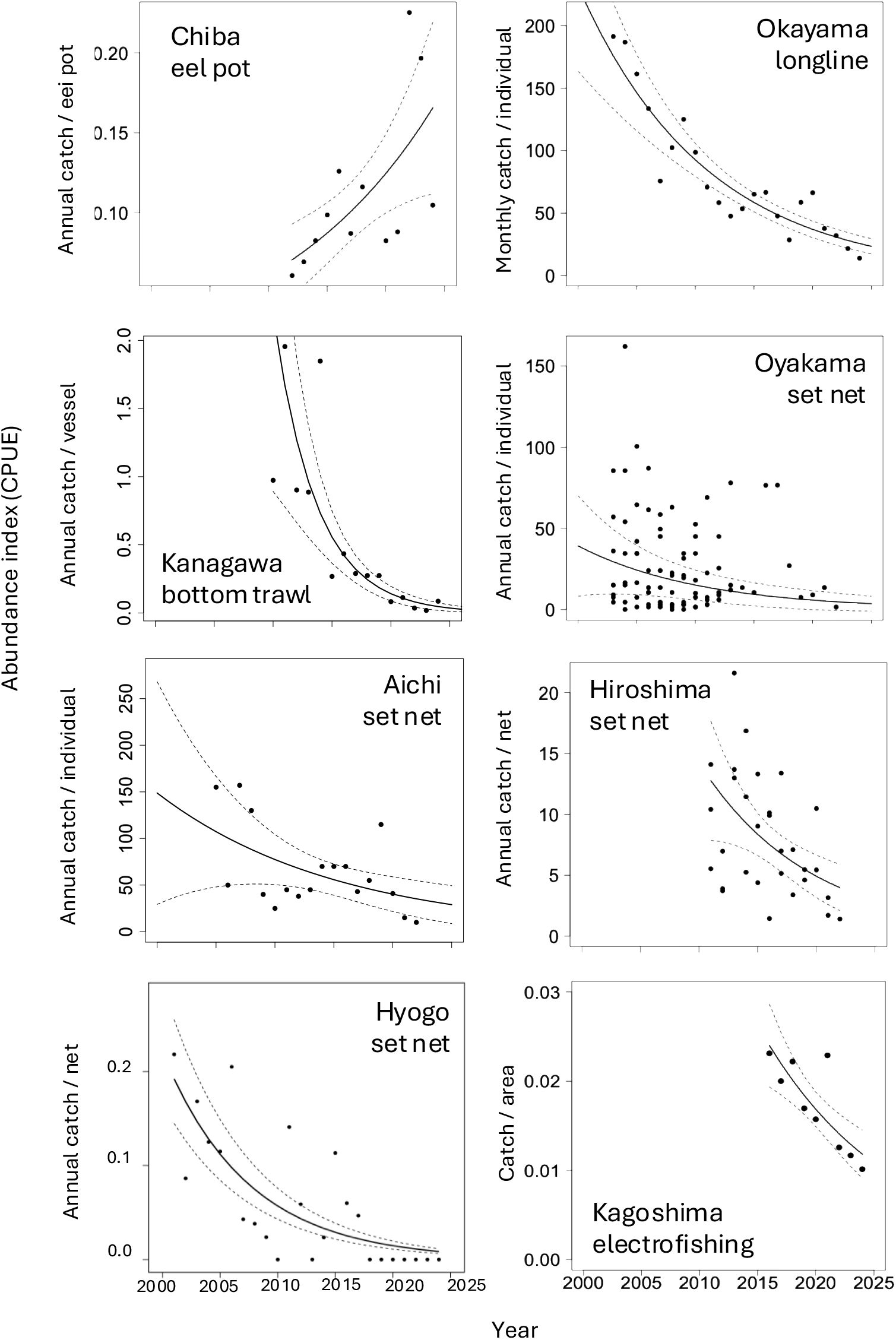
Abundance indices (CPUE) of yellow and silver eels in Japan. The plots indicate observed CPUEs in each year. The solid and dotted lines indicate estimated CPUE decrease/increase and 95% confidential intervals. Please refer to Table 1-3 for details.

For glass eels, this study used data from two scientific monitoring programmes. The first is the data from the ‘Eel River Project’ conducted in the Sagami River, Japan, where monthly surveys have been carried out during spring tide periods since 2009 (Aoyama et al. 2012). The second dataset is from the Yilan River in Taiwan, where collaborative monitoring between fishers and researchers has been ongoing since 2012. Information about both monitoring programmes is provided in Table 1. For the Sagami River, scoop nets were used to capture glass eels. Monitoring of glass eels was conducted at night during spring tides or on nights close to the spring tide, specifically during the flood tide between low and high tide. In the Sagami River, monitoring involved the use of lights to visually locate and scoop up incoming glass eels. Further details are provided in Aoyama et al. (2012). In the Yilan River estuary, researchers record the catch and number of nets used in glass eel fishing conducted by local fishers who used small, fixed fyke nets. The glass eels were collected once or twice a month at night during flood tides between 2012 and 2025. For Japanese eels, glass eels migrate to East Asia from around November to May. Since the migration period spans calendar years, catch data were compiled on a yearly basis from November of the previous year to October of the current year, following the practice of administrative agencies in East Asia (Fisheries Management and Scientific Research Departments of the People’s Republic of China et al. 2025).

FW in the ‘Environment’ column indicates freshwater environments, while BW denotes brackish water environments. In the ‘Fishery dependence’ column, ‘Fishery’ refers to data derived from fishery-dependent sources, whereas ‘Independent’ indicates data obtained through scientific monitoring independent from commercial fisheries. ^*^: fishery data was not recorded in 2019. ^**^: glass eel monitoring at Sagami River was suspended between 2020 and 2021 due to restrictions on movement associated with the COVID-19 pandemic.

### Statistical analysis

We analysed the relationship between CPUE and year using generalised linear models (GLMs) or generalised linear mixed models (GLMMs) (Table 2). The response variable was catch, expressed either as biomass (kg) or as number of individuals, depending on the dataset. Given the frequent occurrence of low catch values, the gamma distribution was selected as the probability distribution when the catch was reported as weight, whereas a negative binomial distribution was applied when the catch was recorded as the number of individual eels (yellow and silver eel data from Chiba and Kagoshima and glass eel data). Among the datasets analysed in this study, only the one from Hyogo included records with zero catch. Although set-net fishing has continued in this water body, no eel catches have been recorded at all since 2018. Since the gamma distribution cannot be applied to data that include zeros, a Tweedie distribution was selected for this dataset. To analyse CPUE, fishing effort, such as number of gears or fishing duration, was included as an offset term, and a log link function was applied. However, when only one fisher was involved and no other information on effort was available, no offset term was included. Where GLMMs were applied, individual fishers were treated as random effects. Differences in models, variables, offset terms, and random effects for each dataset are summarised in Table 2.

**Table 2.** Models and variables used in analysis on eel abundance indices.

| Life stage | Location | Model | Response variable | Explanatory variable | Offset term | Random effect |
| --- | --- | --- | --- | --- | --- | --- |
| Yellow & silver eel | Chiba | GLM | Number | Year | Eel pots | NA |
|  | Kanagawa | GLM | Weight | Year | Vessels | NA |
|  | Aichi | GLM | Weight | Year | none | NA |
|  | Hyogo | GLM | Weight | Year | Nets | NA |
|  | Okayama longline | GLM | Weight | Year | Month of fishing | NA |
|  | Okayama set net | GLMM | Weight | Year | none | Individual fisherman |
|  | Hiroshima | GLMM | Weight | Year | Nets | Individual fisherman |
|  | Kagoshima | GLM | Number | Year | Surface area | NA |
| Glass eel | Sagami River | GLM | Number | Year | Hours × nets | NA |
|  | Yilan River | GLM | Number | Year | Hours × nets | NA |

Based on the coefficients obtained from the analyses, we calculated the rate of decline over a three generation length. This period, 24 years in the case of Japanese eels, is used in Criterion A of the IUCN Red List assessments to evaluate population trends (Pike et al. 2020a). At the IUCN, which manages the Red List, population size is measured only in terms of mature individuals (IUCN 2024). In the case of freshwater eel species, it is extremely difficult to estimate the number of spawners that actually reach the spawning areas. Instead of fully matured adults, the number of silver eels undertaking downstream migration for spawning is widely used as a proxy indicator for the number of mature individuals (e.g., Pike et al. 2020b). Because Japanese eels undergo their spawning migration from fall to early winter (Sudo et al. 2017), eel catches from October to December can contain a large proportion of silver eels. Therefore, following Kaifu et al. (2018), we estimated the proportion of eel catches during this season relative to the total annual catch.

All statistical analyses were performed using R version 4.4.1 (R Core Team 2024). The following R packages were used: ‘glmmTMB’ for fitting GLMMs and ‘MASS’ for model fitting and diagnostics.

The offset term was based on the number of elements indicated in the ‘Offset term’ column. For example, in the case of ‘Okayama longline’, where fisheries data of a single fisher was obtained, the offset term corresponds to the number of months during which fishing was conducted. When multiple fishers were involved and individual catch data were available, the individual fisher was treated as a random effect, and the number of fishing gear operated by that fisher was used as the offset term. For example, in ‘Okayama set net’, since the number of nets used by each fisher was unknown, no offset term was applied. For glass eel data, the offset term was defined as the fishing duration (hours) and the number of nets.

## Results

Of the eight datasets on CPUE for yellow and silver eels, seven showed statistically significant declines, while the remaining one exhibited a significant increase (Table 3, Figure 2). Among the seven datasets with significant declines, the projected decrease over a three-generation period (24 years) ranged from 79.2% to 99.9%. In Chiba, where a significant increase was detected, the model suggested an approximate 4.5-fold increase over the same time period. The proportion of total annual catch occurring during the autumn to early winter period (October to December) varied substantially among datasets, ranging from 9.3% in Chiba to 92.4% in Kanagawa (Table 3). In Kanagawa, where most eel catches are concentrated from autumn to early winter, a sharp decline in catch was observed (Fig. 4).

**Table 3.** Results of abundance index (CPUE) analysis for yellow and silver eels in Japan.

| Location |  | Estimate | SE | t/z value | p value | Depletion rate (%) | % catch in Oct-Dec |
| --- | --- | --- | --- | --- | --- | --- | --- |
| Chiba | Intercept | -146.832 | 40.939 | -3.587 | < 0.001 | -452.2 | 9.3 |
|  | Coefficient | 0.0717 | 0.020 | 3.532 | < 0.001 |  |  |
| Kanagawa | Intercept | 553.065 | 74.352 | 7.438 | < 0.0001 | 99.9 | 92.4 |
|  | Coefficient | -0.291 | 0.037 | -7.778 | < 0.0001 |  |  |
| Aichi | Intercept | 135.713 | 57.032 | 2.380 | < 0.05 | 79.2 | Unknown |
|  | Coefficient | -0.065 | 0.028 | -2.307 | < 0.05 |  |  |
| Hyogo | Intercept | 268.758 | 67.987 | 3.953 | < 0.0001 | 96.1 | Unknown |
|  | Coefficient | -0.135 | 0.034 | -3.989 | < 0.0001 |  |  |
| Okayama longline | Intercept | 188.893 | 20.199 | 9.352 | < 0.0001 | 88.9 | 16.1 |
|  | Coefficient | -0.092 | 0.010 | -9.143 | < 0.0001 |  |  |
| Okayama set net | Intercept | 2.850 | 0.306 | 9.330 | < 0.0001 | 89.9 | Unknown |
|  | Coefficient | -0.096 | 0.034 | -2.814 | < 0.01 |  |  |
| Hiroshima | Intercept | 2.045 | 0.103 | 19.925 | < 0.0001 | 92.2 | 68.2 |
|  | Coefficient | -0.106 | 0.035 | -3.018 | < 0.01 |  |  |
| Kagoshima | Intercept | 175.064 | 45.708 | 3.830 | < 0.001 | 88.1 | NA |
|  | Coefficient | -0.089 | 0.023 | -3.918 | < 0.0001 |  |  |

The ‘Depletion rate’ column indicates the estimated decline over a 24-year period— equivalent to the three-generation length of *Anguilla japonica*—based on the model estimations. The negative value for Chiba indicates an increase. The ‘% catch in Oct–Dec’ column shows the proportion of catch occurring between October and December comparing to the annual catch, which corresponds to the downstream spawning migration period of this species. In the case of ‘Kagoshima’, visual confirmation by experts was made during sampling to ensure that no silver eels were included. Monthly catch data were not available for ‘Aichi,’ ‘Hyogo,’ and ‘Okayama set net’.

For glass eels, no significant correlation was detected between CPUE and year (Table 4). CPUE exhibited substantial monthly and annual variability, and time-series plot of CPUE did not suggest a declining trend (Figure 3).

**Table 4.** Results of CPUE analysis for glass eels.

| Location | Fishing method |  | Estimate | SE | z value | p value |
| --- | --- | --- | --- | --- | --- | --- |
| Sagami River | Scoop net | Intercept | 44.715 | 63.026 | 0.709 | 0.478 |
|  |  | Coefficien | -0.021 | 0.0313 | -0.678 | 0.498 |
| Yilan River | Set net | Intercept | -200.510 | 121.151 | -1.655 | 0.098 |
|  |  | Coefficien | 0.101 | 0.060 | 1.675 | 0.094 |

**Fig. 3.**
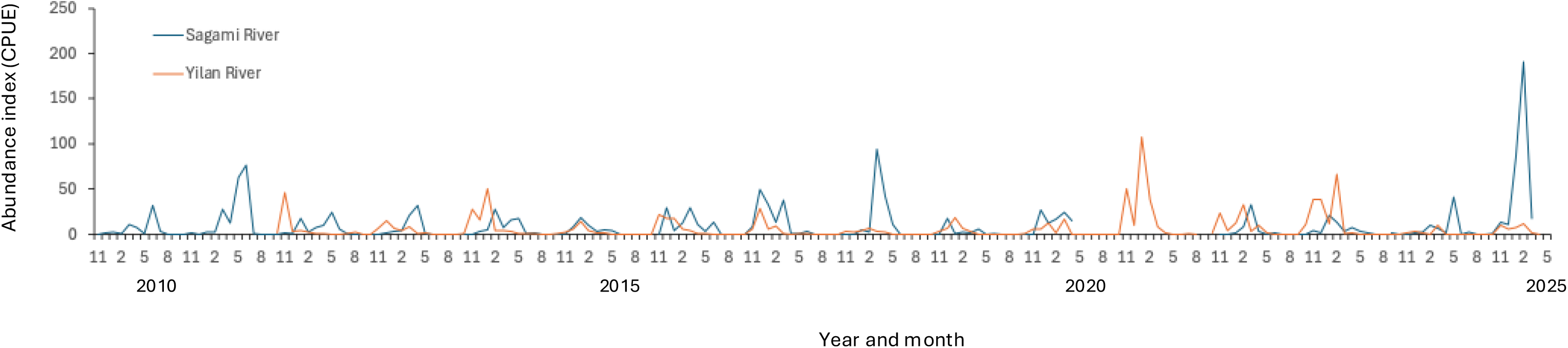
Abundance indices (CPUE) of glass eels collected in Japan and Taiwan For the Sagami River, Japan, CPUE is shown as the number of individuals per unit of time and number of nets, based on monitoring using scoop nets. For the Yilan River, Taiwan, researchers collected fishery data from small set nets, and CPUE is presented as the number of individuals per unit of time and number of nets. Although glass eels of *Anguilla marmorata* also migrate to Japan and Taiwan, only Japanese eels (*A. japonica*) were selected for the CPUE calculation. Please refer to Table 4 for analytical results on annual trends.

**Fig. 4.**
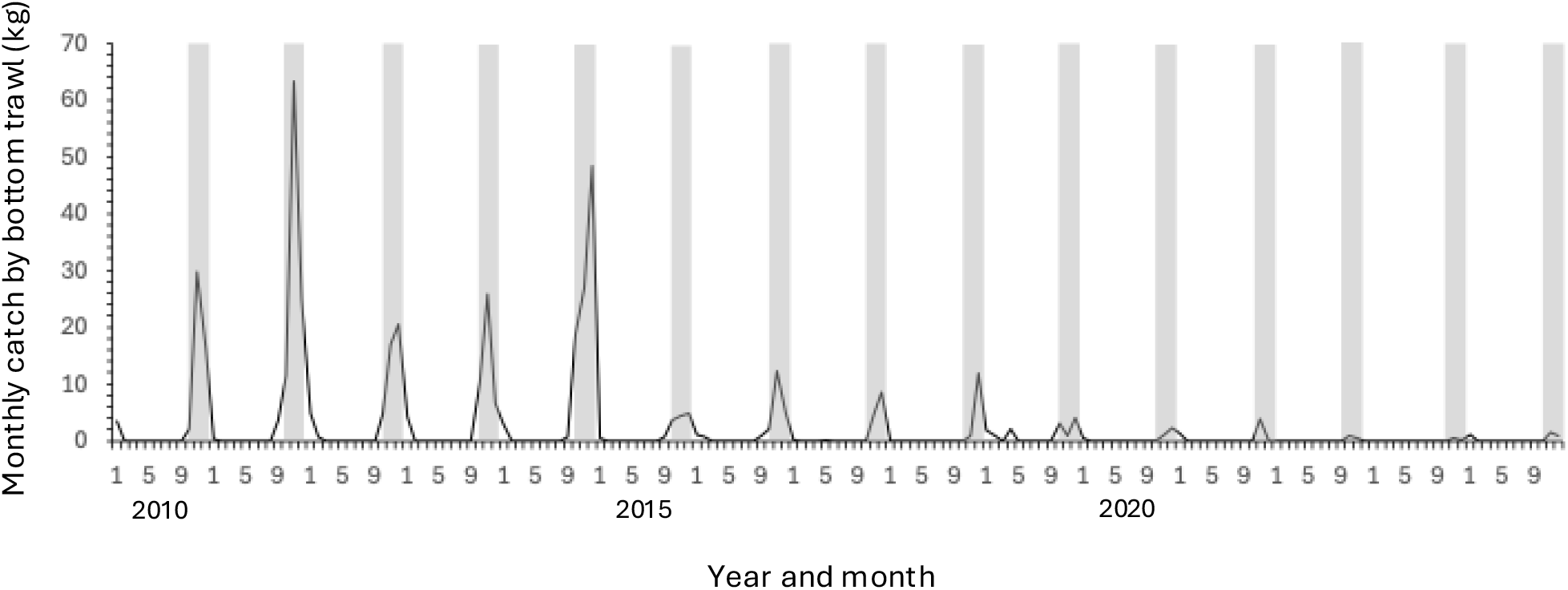
Monthly trends in eel catch by bottom trawl fisheries in Kanagawa Prefecture Periods between October and December are shaded.

In all seven fishery-related datasets, the number of fishers remained constant or declined during the study period. In Kanagawa, the number of vessels operating bottom trawl fisheries declined from 53 in 2010 to 43 in 2024. In Hyogo, the set-net fishery had seven fishers and 22 nets in 2001, but this had decreased to five fishers and 11 nets by 2024. Furthermore, in three water bodies, Aichi, Okayama set net, and Hiroshima, the targeted fisheries had ceased entirely. Given that fishing pressure in the locations where yellow and silver eel fisheries data were collected has remained stable or declined, overall fishing pressure on Japanese eels is considered to have decreased, at least the datasets used in this study. These reductions in fishing pressure were accounted for in the model analyses (see Materials and Methods for details).

## Discussion

For yellow and silver eels, statistically significant declines in CPUE were detected in seven of the eight datasets, while a significant increase was observed in one. Taken together, the overall trend across all datasets indicates a decline in eel abundance, although increases were observed in certain areas. Notably, the decline in CPUE in Kanagawa, where 92.4% of the total annual catch occurred during the downstream spawning migration period, is of particular concern. In this fishery, eels are not the primary target but are caught as bycatch, with white spotted conger (*Conger myriaster*) being one of the main target species. Given that almost no eels are caught in spring and summer, it can be inferred that few eels inhabit this area year-round, and that silver eels migrating towards the spawning grounds are being caught from autumn to winter. In Red List assessments by the IUCN, the number of silver eels is considered the most appropriate indicator of population trends in anguillid eels (Pike et al. 2020b). In the present study, the CPUE of eel catches in Kanagawa—presumed to consist primarily of silver eels—was estimated to have declined by 99.9% over a three-generation length (24 years) (Pike et al. 2020a). Similarly, in Hiroshima, where the proportion of catches during the downstream migration season was relatively high at 68.2%, CPUE also showed a significant decline, with a projected decrease of 92.2% during three-generation length. In contrast, the only dataset showing an increase in CPUE was from Chiba, where the proportion of catches during the migration season was low at 9.3%, suggesting that most of the eels caught there were yellow eels.

In contrast to the results for yellow and silver eels, no significant correlation was detected between CPUE and year in the two datasets for glass eels. The recruitment of glass eels is strongly influenced by oceanic conditions, such as ocean currents, and is subject to substantial interannual fluctuations (Kimura and Tsukamoto 2006). While the volume of glass eel recruitment may potentially serve as a long-term indicator of population dynamics, clear trend was not detected in the relatively short timescale of approximately 15 years analysed in this study.

In this study, the six eel fishery datasets reported by Kaifu and Yokouchi (2019) were updated, and two additional datasets from freshwater environments were incorporated. As the abundance indices (CPUE) for yellow and silver eels were derived from areas where the proportion of stocked individuals is low, the influence of stocking on the data is expected to be minimal. For glass eels, rather than relying on official fishery statistics—which are considered unreliable due to the presence of unreported catch—this study utilised scientific monitoring data. Through these measures, the data provided by this study provided the best currently available indicators of population dynamics of the Japanese eel.

## Acknowledgements

We express our sincere gratitude to the fishers and fisheries cooperative associations who kindly provided fisheries data for this study. We also appreciate Drs. J. Aoyama, N. Amiya, F. Furukawa, and S. Tsutsui for their help with the scientific monitoring of glass eel. This study was financially supported by the Asahi Glass Foundation, Chuo University, Japan Fisheries Agency (Eel Supply Stabilization Project), JSPS KAKENHI (16K07851, 21K05751, 22H00371), National Science and Technology Council, Executive Yuan, Taiwan (MOST 111-2313-B-002-016-MY3) (alphabetical order).

## References

Aoyama, J., Miller, M.J., 2003. The silver eel. In: Aida, K., Tsukamoto, K., Yamauchi, K. (Eds.), Eel Biology. Springer, Tokyo, pp. 107–117. 10.1007/978-4-431-65907-5_8

Aoyama, J., Shinoda, A., Yoshinaga, T., Tsukamoto, K., 2012. Late arrival of Anguilla japonica glass eels at the Sagami River estuary in two recent consecutive year classes: ecology and socio-economic impacts. Fish. Sci. 78, 1195–1204. 10.1007/s12562-012-0544-y

Fisheries Management and Scientific Research Departments of the People’s Republic of China, the Fisheries Agency of Japan, the Ministry of Oceans and Fisheries of the Republic of Korea and the Fisheries Agency of Chinese (2025) Joint Press Release. https://www.maff.go.jp/j/pr/event/attach/pdf/kaigi.release-1625.pdf

Itakura, H., Arai, K., Kaifu, K., Shirai, K., Yoneta, A., Miyake, Y., Secor, D.H., Kimura, S., 2018. Distribution of wild and stocked Japanese eels in the Tone River watershed revealed by otolith stable isotopic ratios. J. Fish Biol. 93, 805–813. 10.1111/jfb.13782

IUCN Standards and Petitions Committee, 2024. Guidelines for Using the IUCN Red List Categories and Criteria. Version 16. IUCN, Gland, Switzerland. https://nc.iucnredlist.org/redlist/content/attachment_files/RedListGuidelines.pdf

Jacoby, D., Gollock, M., 2014. Anguilla japonica. The IUCN Red List of Threatened Species. Version 2014.3. https://www.iucnredlist.org/species/166184/1117791

Kaifu, K., 2019. Challenges in assessments of Japanese eel stock. Mar. Policy 102, 1–4. 10.1016/j.marpol.2019.02.005

Kaifu, K., Yokouchi, K., Higuchi, T., Itakura, H., Shirai, K., 2018a. Depletion of naturally recruited wild Japanese eels in Okayama, Japan, revealed by otolith stable isotope ratios and abundance indices. Fish. Sci. 84, 757–763. 10.1007/s12562-018-1225-2

Kaifu, K., Yokouchi, K., 2019. Increasing or decreasing? – current status of the Japanese eel stock. Fish. Res. 220, 105348. 10.1016/j.fishres.2019.105348

Kimura, S., Tsukamoto, K., 2006. The salinity front in the North Equatorial Current: a landmark for the spawning migration of the Japanese eel (Anguilla japonica) related to the stock recruitment. Deep Sea Res. II 53, 315–325. 10.1016/j.dsr2.2006.01.009

Pike, C., Kaifu, K., Crook, V., Jacoby, D., Gollock, M., 2020a. Anguilla japonica (amended version of 2020 assessment). The IUCN Red List of Threatened Species 2020: e.T166184A176493270. 10.2305/IUCN.UK.2020-3.RLTS.T166184A176493270.en

Pike, C., Crook, V., Gollock, M., 2020b. Anguilla anguilla. The IUCN Red List of Threatened Species 2020: e.T60344A152845178. 10.2305/IUCN.UK.2020-2.RLTS.T60344A152845178.en

R Core Team, 2024. R: A language and environment for statistical computing. R Foundation for Statistical Computing, Vienna, Austria. https://www.R-project.org/

Righton, D., Piper, A., Aarestrup, K., Amilhat, E., Belpaire, C., Casselman, J., Castonguay, M., Diaz, E., Doerner, H., Faliex, B., Feunteun, E., Fukuda, N., Hanel, R., Hanzen, C., Jellyman, D., Kaifu, K., McCarthy, K., Miller, M., Pratt, T., Sasal, P., Schabetsberger, R., Shiraishi, H., Simon, G., Sjoberg, N., Steele, K., Tsukamoto, K., Walker, A., Westerberg, H., Yokouchi, K., Gollock, M., 2021. Important questions to progress science and sustainable management of anguillid eels. Fish. Fish. 22, 762–788. 10.1111/faf.12549

Shiraishi, H., Crook, V., 2015. Eel market dynamics: an analysis of Anguilla production, trade and consumption in East Asia. TRAFFIC, Tokyo. https://www.traffic.org/publications/reports/eel-market-dynamics-an-analysis-of-anguilla-production-trade-and-consumption-in-east-asia/

Sudo, R., Okamura, A., Fukuda, N., Miller, M.J., Tsukamoto, K., 2017. Environmental factors affecting the onset of spawning migrations of Japanese eels (Anguilla japonica) in Mikawa Bay, Japan. Environ. Biol. Fish. 100, 237–249. 10.1007/s10641-017-0575-4

Tanaka, E., 2014. Stock assessment of Japanese eels using Japanese abundance indices. Fish. Sci. 80, 1129–1144. 10.1007/s12562-014-0807-x

Tsukamoto, K., 1990. Recruitment mechanism of the eel, Anguilla japonica, to the Japanese coast. J. Fish. Biol. 36, 659–671. 10.1111/j.1095-8649.1990.tb04320.x

Tsukamoto, K., Chow, S., Otake, T., Kurogi, H., Mochioka, N., Miller, M.J., Aoyama, J., Kimura, S., Watanabe, S., Yoshinaga, T., Shinoda, A., Kuroki, M., Oya, M., Watanabe, T., Hata, K., Ijiri, S., Kazeto, Y., Nomura, K., Tanaka, H., 2011. Oceanic spawning ecology of freshwater eels in the western North Pacific. Nat. Commun. 2, 1–9. https://www.nature.com/articles/ncomms1174

Wakiya, R., Itakura, H., Hirae, T., Igari, T., Manabe, M., Matsuya, N., Miyata, K., Sakata, M.K., Minamoto, T., Yada, T., Kaifu, K., 2022. Slower growth of farmed eels stocked into rivers with higher wild eel density. J. Fish. Biol. 101, 613–627. 10.1111/jfb.15131

